# Deficient skeletal muscle regeneration after injury induced by a *Clostridium perfringens* strain associated with gas gangrene

**DOI:** 10.1101/441204

**Authors:** Ana Mariel Zúñiga-Pereira, Carlos Carlos Santamaría, José María Gutiérrez, Alberto Alape-Girón, Marietta Flores-Díaz

## Abstract

Very little is known about the muscle regeneration process that follows myonecrosis induced by *C. perfringens*, the main agent of gas gangrene. This study revealed that, in a murine model of the infection with a sublethal inoculum of *C. perfringens*, muscle necrosis occurs concomitantly with significant vascular damage, which limits the migration of inflammatory cells. A significant increase in cytokines that promote inflammation explains the presence of inflammatory infiltrate; however, an impaired IFNγ expression, a reduced number of Ml macrophages, a deficient phagocytic activity, and the prolongation of the permanence of inflammatory cells, lead to deficient muscle regeneration. The expression of TGFβ1 and the consequent accumulation of collagen in the muscle, likely contribute to the fibrosis observed 30 days after infection. These results provide new information on the pathogenesis of gas gangrene caused by *C. perfringens*, shed light on the basis of the poor muscle regenerative activity, and may open new perspectives for the development of novel therapies for patients suffering this disease.

## Introduction

Muscle regeneration after myonecrosis occurs in three sequential and interrelated phases: inflammation, regeneration, and remodeling (Carosio et al., 2011). Initially, cellular damage is associated with the entry of extracellular calcium that induces a series of degenerative events, including hypercontraction, mitochondrial alterations and the activation of calcium-dependent proteases, leading to necrosis of the myofibers (Carosio et al., 2011; Turner and Badylak, 2012). Moreover, the disruption of the sarcolemma results in an increase in serum levels of creatine kinase (CK), a protein normally restricted to the myofiber cytosol (Karalaki et al. 2009). The presence of necrotic fibers activates the inflammatory response and then an influx of specific cells of the immune system occurs in the damaged muscle (Carosio et al., 2011). Inflammation is a critical component of the regenerative process (Carosio et al., 2011). Injured muscle fibers activate the synthesis and release of a plethora of signaling molecules into the extracellular space, and these mediators induce the sequential attraction and activation of diverse cell populations that promote muscle regeneration (Tidball, 2011; Karalaki et al., 2009). The vascular network has an important role in skeletal muscle regeneration as it has an impact on the distribution of recruited inflammatory cells, regeneration-related factors (growth factors, cytokines, chemokines), as well as nutrients. Therefore alterations in vascular integrity can affect the regenerative process (Gutiérrez et al., 2018).

Muscle regeneration begins with the activation of satellite cells (SC) that reside on the surface of muscle fibers (Tidball, 2017). Following muscle damage, some SC proliferate and differentiate whereas others return to quiescence as reserve population of myogenic cells (Tidball, 2017). Postmitotic precursor cells derived from activated SC then form multinucleated myotubes and proceed through a stage of regeneration that is dominated by terminal differentiation and growth (Tidball, 2017). When the formation of contractile muscle fibers is complete, the size of the newly formed myofibers increases and the nucleus is displaced to the periphery of the fiber (Karalaki et al., 2009).

Remodeling is a process in which the maturation and functional performance of regenerated myofibers occurs (Carosio et al., 2011). The final phase of the regenerative process is characterized by the remodeling of connective tissue, angiogenesis, and functional recovery of injured skeletal muscle (Carosio et al., 2011). After muscle injury, the extracellular matrix is remodeled, resulting in the overproduction of several types of collagen that contribute to the formation of a scar (Carosio et al., 2011); however, the overproduction of collagens within the damaged area could lead to excessive scarring and loss of muscle function (Carosio et al., 2011). The transforming growth factor β1 (TGFβ1) has been identified as a key factor in the activation of the fibrosis cascade in injured skeletal muscle (Carosio et al., 2011). The processes of neovascularization and reinnervation play a critical role in determining the regeneration potential of the injured muscle (Turner and Badylak, 2012).

The influx of inflammatory cells to the site of muscle damage is paramount for efficient regeneration (Chazaud et al., 2003). The inflammatory response during the early stages of muscle regeneration is temporally and spatially coupled to the initial stages of myogenesis, when SC are activated and initiate their proliferation and differentiation (Tidball, 2017). Polymorphonuclear leucocytes (PMN) (Ly6C^+^) are the first inflammatory cells to invade the damaged muscle. The intramuscular density of these cells increases on the first six hours after the muscular damage, reaching a peak 24 h after the injury and then gradually returning to normal (Karalaki et al., 2009; Tidball, 2011; Tidball, 2017). PMN are essential for microbicidal action and for attracting other leucocytes capable of resolving inflammation and mediating the regeneration process (De Filippo et al., 2008). Resident tissue macrophages (F4/80^+^, LY 6C^+^) promote a marked flow of PMN through the release of the main chemoattractants for the recruitment of PMN, such as the murine chemokine keratinocyte chemo attractant (KC or CXCL1) and macrophage inflammatory protein 2 (MIP2 or CXCL2). PMN initiate the process of removal of necrotic myofibers and cellular debris by phagocytosis and by the rapid release of high concentrations of free radicals and proteases (Turner and Badylack, 2012; Carosio et al., 2011). In addition, PMN secrete proinflammatory cytokines that stimulate the arrival of macrophages, further promoting tissue inflammation (Turner and Badylack, 2012; Carosio et al., 2011).

Two distinct populations of macrophages sequentially invade the damaged muscle tissue. A population of phagocytic macrophages CD68^+^/ CD163^−^/F4/80^+^ (M1 macrophages) closely follow the invasion of PMN, reaching their maximum concentration approximately two days after the injury, and subsequently decreasing in number (Tidball, 2011). The Th1 response is characterized by the presence of interleukin-1 (IL1β), interleukin-2 (IL2), interferon-γ (IFNγ) and tumor necrosis factor alpha (TNFα) (Tidball, 2008). In turn, the M1 macrophages that produce TNFα and IL1β are capable of damaging the host tissue by releasing oxygen-free radicals that can damage cell membranes, phagocytose the necrotic muscle and promote the proliferation of SCs (Carosio et al., 2011; Turner and Badylack, 2012). The Th2 response involves high levels of interleukin-4 (IL4), interleukin-5 (IL5), interleukin-6 (IL6), interleukin-10 (IL10) and interleukin-13 (IL13), which have anti-inflammatory effects and deactivate M1 macrophages (Tidball, 2008). Additionally, IL4, IL10, and IL13 play well-characterized roles in the activation of non-phagocytic macrophages (Tidball, 2008; Tidball and Villalta, 2010). The non-phagocytic macrophages CD68^−^/CD163^+^/F480^+^/206^+^, known as M2 macrophages, invade the muscle and reach their maximum peak approximately four days after injury, but the number of these cells remains elevated in the damaged muscle by periods of up to two weeks (Tidball, 2011). M2 macrophages express high levels of IL10 and low levels of IL12 (Carosio et al., 2011). These tissue-remodeling macrophages decrease the inflammatory response and promote angiogenesis, as well as myoblast proliferation, growth, and differentiation (Carosio et al., 2011; Turner and Badylack, 2012).

Stimulation by IFNγ is essential for the classic activation of the Th1 phenotype (Tidball, 2008). IFNγ is also a powerful activator of PMN and M1 macrophages (Tidball, 2017). Although IFNγ is usually a product of "natural killer" cells and T cells, it could also be expressed by M1 macrophages having an autocrine role in their activation (Tidball, 2011; Tidball, 2017). Furthermore, the stimulation of IFNγ can increase the response of PMN to chemotactic cytokines, potentially increasing their invasion to the sites of injury (Tidball, 2011).

TNFα is another Th1 cytokine highly expressed by M1 macrophages. TNFα in the muscle reaches its peak approximately 24 h after the onset of damage, which coincides with the invasion of PMN and M1 macrophages, and with the increase in secondary muscle damage generated by myeloid cells (Tidball, 2011). Part of myeloid cell-mediated damage to muscle fibers is caused by nitric oxide (NO) derived from inducible nitric oxide synthase (iNOS) and TNFα can stimulate M1 macrophages to elevate iNOS expression and thus promote further damage to muscle fibers (Tidball, 2011). The potential of TNFα to promote muscle repair and regeneration lies in its direct action on muscle cells (Tidball, 2017).

Another important factor that influences the muscle regeneration process is TGFβ1, which has been recognized as a modulator of myoblast activity (Chargé and Rudnicki, 2004). In general, TGFβ1 plays a negative role in the regulation of myogenesis; it is highly expressed in quiescent SC and represses the progress of the cell cycle in these cells repressing the expression of MyoD and myogenin. (Fu et al., 2015).

Gas gangrene induced by *Clostridium perfringens* is an acute and life threatening infection associated either to trauma or surgery and is characterized by fever, sudden onset of prominent pain, the accumulation of gas at the site of infection, massive local edema and a severe myonecrosis (Stevens and Bryant, 2017). When there is an anaerobic environment adequate for clostridial growth after the introduction of *C. perfringens* in a deep lesion or in a surgical wound, bacteria begin to multiply and the destruction of the muscle spreads within few hours (Stevens and Bryant, 2017). In mice, intramuscular injection of 10^6^ wild type *C. perfringens* vegetative cells leads to a limited sublethal infection characterized by swelling and myonecrosis. In this work this model was used to characterize the regeneration process after skeletal muscle damage induced by this bacterium.

## Materials and Methods

### Bacterial culture

*C. perfringens* (strain JIR325) was grown on Brain Heart Infusion (BHI) broth in an anaerobic chamber until an OD_600_ of 0.47 was reached. The number of colony forming units (CFU) per 100 μl was determined by plating serial 10-fold dilutions on BHI agar plates supplemented with yolk.

### Experimental infection

CD-1 mice of 18-20 g body weight were injected in the left gastrocnemius with 1×10^6^ CFU of *C. perfringens* JIR325, in 100 μl of 0.12 M NaCl, 0.04 M phosphates, pH 7.2 (PBS). All the procedures involving the use of animals in this study were approved by the Institutional Committee for the Care and Use of Laboratory Animals (CICUA) of Universidad de Costa Rica (approval number CICUA-098-17), and meet the Animal Research Reporting in vivo Experiments (ARRIVE) guidelines, and the International Guiding Principles for Biomedical Research Involving Animals of the Council of International Organizations of Medical Sciences (CIOMS).

### CK activity assay

To evaluate myotoxicity, blood samples were collected 5 and 24 h post infection, and the CK activity in plasma was determined using the “CK-NAC UV Unitest” (Wiener Lab, Argentina) according to the manufacturer’s instructions.

### Histological analysis

For histological analysis, groups of 3 mice were injected with 1×10^6^ CFU of *C. perfringens* JIR325 or with sterile PBS. Animals were sacrificed in the phase of muscle damage at 1, 5, and 24 h post infection, and in times that cover the various steps in the process of muscle regeneration, i.e., 3, 5, 7, 14 and 30 d post infection. The injected gastrocnemius muscles were dissected out and placed in a zinc fixative solution (calcium acetate 3 mM, zinc acetate 27 mM, zinc chloride 36 mM, Tris buffer 0.1 M, pH 7.4), for at least 48 h at 4°C. The dissected muscles were dehydrated in ethanol, placed in xylene, and embedded in paraffin. Three non-consecutive sections of 4 μm were obtained from the mid region of each muscle and placed in glass slides. Sections were deparaffinated in xylene, hydrated in distilled water and stained with hematoxylin and eosin. The microscopic evaluation was performed in an OLYMPUS BX51 microscope. Images of total muscle were captured from each section using an Evolution MP camera (Media Cybernetics, USA) and analyzed using the image analysis software Image Pro 6.3 (Media Cybernetics, USA). The necrotic area was estimated in samples collected 24 h post infection, considering the percentage of the area observed corresponding to damage and hypercontracted fibers. Areas of regeneration and of lack of regeneration were estimated in samples collected 14, and 30 d post infection; areas of regeneration corresponded to the percentage of the examined area characterized by the presence of regenerating fibers (fibers with centrally located nuclei), while non-regenerative areas were defined as the percentage of the examined area corresponding to cell debris and fibrotic muscle, while The diameters of regenerating fibers were determined in sections of muscles collected 30 d post infection.

### Collagen Staining

Groups of 3 mice were injected with *C. perfringens* JIR325 or with sterile PBS, and sacrificed 7, 14, and 30 d post infection. Increments of collagen in the muscle were detected by staining with Direct Red 80 (Sigma-Aldrich, USA) (0.1% in a saturated picric acid solution) which stains collagen, and Fast Green FCF 0.1% (Sigma-Aldrich, USA), which stains other proteins, for one h at room temperature, according with the procedure described by Hernández *et al.* (2011). Slides were washed with acidified water (5 mL of glacial acetic acid per liter), dehydrated, and cleared in xylene. Microscopic evaluation was performed in an OLYMPUS BX51 microscope. Images of total muscle were captured from each section using an Evolution MP camera (Media Cybernetics, USA) and analyzed using the image analysis software Image Pro 6.3 (Media Cybernetics, USA). The percentage of fibrosis (collagen deposition) in total muscle was quantified 30 d post infection, using the image analysis software ImageJ 1.51K (National Institutes of Health, USA).

### Quantitative PCR

Relative expression of transcripts coding for IL1β, IL6, TNFβ_1_, INFγ, TGFγ, IL13, IL10, MIP2, KC and MCP-1 was determined in a similar schedule as reported in other muscle injuries models (Tidball, 2008, Tidball, 2011, Tidball, 2017). Groups of 6 mice were injected with *C. perfringens* JIR325 or with sterile PBS in the left gastrocnemius, and sacrificed 6 h, 1, 2, 3 and 6 d post-infection. The left gastrocnemius muscles were rapidly dissected out and ground under sterile conditions. Total RNA was extracted using TRIzol^®^Reagent (ambion, Invitrogen), according to the manufacturer’s instructions and quantified using a NanoDrop 2000c Spectrophotometer (Thermo Scientific, USA) RNA was retrotranscripted to cDNA with RevertAid H Minus First Strand cDNA Synthesis Kit (Fermentas, Thermo Fisher Scientific) in a 2720 Thermal Cycler (Applied Biosystems) using 4 μg of total RNA and random hexamers primers. 160 ng of cDNA were used for each reaction in the quantitative real-time PCR (qPCR) using LightCycler^®^ 480 SYBR Green I Master (Roche Diagnostics) and LightCycler^®^ 480 real-time PCR device (Roche Diagnostics). The selected primers to assess inflammatory-response-specific genes PCR are listed in Table 1. Genes used as reference genes were the housekeeping genes gliceraldehide-3-phosphate dehydrogenase (GAPDH), RNA-binding protein S1 (RNSP1) and the ribosomal protein L13A (RPL13A) (Piller et al., 2013). The cycle number at which the reaction crossed an arbitrarily placed threshold (Ct) was determined, and the relative expression of each gene regarding the mean expression of control genes was described using the equation 2-ΔΔCt where ΔCt = Ctgene-Ctcontrol genes (mean) and ΔΔCt =ACtmice+*C. perfringens* −ΔCtcontrol mice (mean) (Livak and Schmittgen, 2001). Left gastrocnemius muscles of healthy mice injected with sterile PBS were used as controls.

**Table 1.**
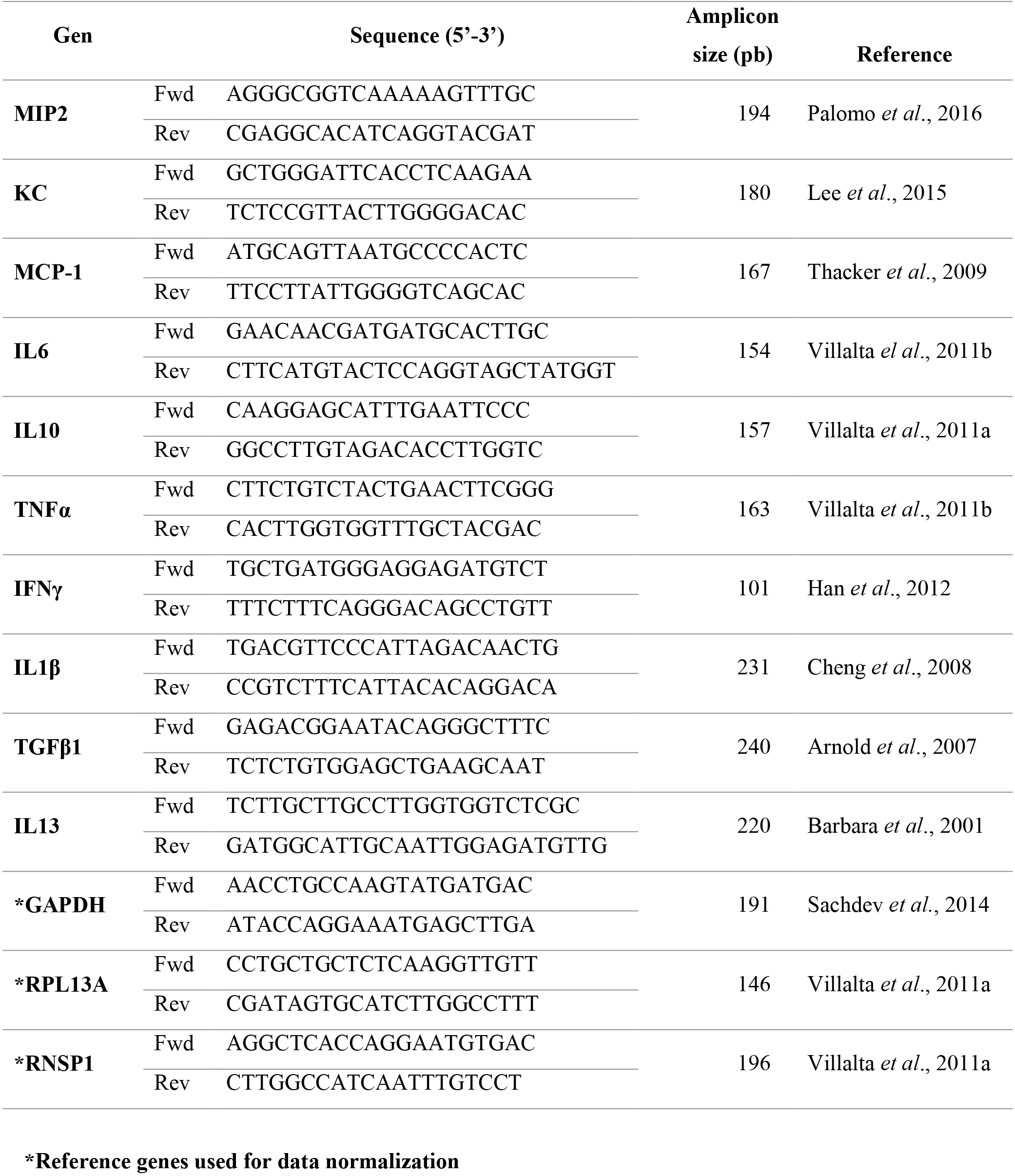
Primers used to assess the expression levels of inflammatory response.

### ELISAs

The IL1β, IL6, TNFα, INFγ, TGFβ1, IL4 and IL10 protein content in the muscle was determined by capture ELISA at times in which their expression had been reported in other muscle injuries (Tidball, 2008, Tidball, 2011, Tidball, 2017). Groups of 6 mice were injected intramuscularly in the left gastrocnemius with *C. perfringens* JIR325 or with sterile PBS. At various time intervals (6 h, 1, 2, 3 and 6 d), mice were killed and the injected gastrocnemius were dissected out, frozen with liquid nitrogen and homogenized in a sterile pyrogen-free saline solution with Complete EDTA-free proteases inhibitor (Roche Diagnostics). Muscle homogenates were centrifuged and the supernatants were collected and stored at −70°C. IL1β, TNFα, INFγ, IL4 and IL10 were quantitated using ELISA kits of R&B Systems (USA), while IL6 and TGFβ_1_ were quantitated using ELISA kits of eBioscience (San Diego, CA, USA), according to the manufacturer’s instructions. Lef gastrocnemius muscles of healthy mice injected with sterile PBS were used as controls.

### Quantification of inflammatory cells

In order to detect different inflammatory cells, groups of three mice were injected with of *C. perfringens* JIR325 or with sterile PBS (negative controls), and sacrificed during the acute (1, 3, 5, and 24 h post infection) and the chronic phase (3, 5, 7 and 14 d post infection). Injected muscles were dissected out and embedded in paraffin, as described previously. For each muscle, three non-consecutive sections of 4 μm were obtained and were placed in positive Chargéd glass slides, deparaffinated in xylene and hydrated. Fluorescence immunohistochemistry for PMN was performed using Anti-Neutrophil Elastase Rabbit pAb (Cat. No. 481001; EMD Millipore); antigen retrieval was carried out by placing the slides in citrate buffer (pH 6) at 50°C for 10 min and blockage steps were performed with Dako Cytomation Biotin Blocking System (Dako, USA), as well as with Protein Block Serum-Free (Dako, Denmark), following the manufacturer’s instructions. Sections were incubated overnight with anti-neutrophil elastase antibody (diluted 1:10) at 4°C in a wet chamber. Then, sections were washed with PBS and incubated with a Polyclonal Goat Anti-Rabbit Immunoglobulins/Biotinylated (Cat. No. E0432; Dako, Denmark) (diluted 1:200) for one h at room temperature. After washing with PBS, sections were incubated with a Streptavidin Alexa Fluor 488 (Cat. No. S11223; Invitrogen) (diluted 1:200) for 30 min at room temperature. Finally, nuclear staining was performed with bis benzimide Hoechst (Cat. No. 33258; Sigma, USA) in a final concentration of 0.5 μg/mL. For the M1 macrophages immunohistochemical stain, the protocol was similar to the one used for PMN with the following modifications: antigen retrieval was carried out using 0.6 U/mL of Proteinase K (Fermentas, Thermo Fisher Scientific) for 5 min at room temperature and the primary antibody used was Rabbit Polyclonal Anti-iNOS (Cat. No. ab15323; Abcam, USA) (diluted 1:75). M2 macrophages were stained with a Goat Plyclonal Anti-Arginase Antibody (Cat. No. ab60176; Abcam, USA). Briefly, antigen retrieval was carried out using Proteinase K (Fermentas, Thermo Fisher Scientific), blockage step was performed with Protein Block Serum-Free (Dako, Denmark) for 10 min at room temperature and then, sections were incubated overnight with the primary antibody (diluted 1:50) at 4°C in a wet chamber. Donkey F(ab’)_2_ antigoat IgG H&L (Alexa Fluor^®^488) preadsorbed (Cat. No. ab150137; Abcam, USA) (diluted 1:200) was used as a secondary antibody, sections were incubated for 1 h at room temperature and then nuclear staining was performed. All the samples were analyzed in a OLYMPUS BX51 microscope, images of the complete histologic sections were captured using an Evolution MP (Media Cybernetics, USA) camera and the number of cells per mm^2^ of the muscle were determined with the Image Pro 6.3 software (Media Cybernetics, USA).

### Quantification of capillaries in muscle tissue

Groups of three mice were injected with *C. perfringens* JIR325 or with sterile PBS (negative controls), and sacrificed at times covering the degenerative phase (6 and 24 h) and the regenerative phase (30 d). Injected muscles were dissected out and embedded in paraffin, as described previously. From the mid region of each muscle, three non-consecutive sections of 4 μm were obtained and placed in positive Chargéd glass slides, and then sections were deparaffinated in xylene and hydrated. Fluorescence immunohistochemistry was carried out in order to detect capillary vessels. For antigen retrieval, Proteinase K (Fermentas, Thermo Fisher Scientific) was used for 5 min at room temperature and then, blockage steps were performed with H_2_O_2_ (3%) for 30 min, Dako Cytomation Biotin Blocking System (Dako, USA) for 10 min each, and with Protein Block Serum-Free (Dako, Denmark) for 1 h. Sections were incubated overnight with purified anti-mouse Flk-1 (IHC) (Cat. No. 555307; BD PharmigenTM) (diluted 1:50) at 4°C in a wet chamber. Then, sections were washed with PBS and incubated with Polyclonal Goat Anti-Rabbit Immunoglobulins/Biotinylated (Cat. No. E0432; Dako, Denmark) (diluted 1:200) for 1 h at room temperature. In order to amplify the signal, the Biotin-XX Tyramide SuperBoostTMKit (Invitrogen) was used according to manufacturer’s instructions. Streptavidin Alexa Fluor 488 (Cat. No. S11223; Invitrogen) (diluted 1:300) was used as final fluorophore for 30 min at room temperature. Nuclear staining was performed with bis BENZIMIDE Hoechst (Cat. No. 33258; Sigma, USA) in a final concentration of 0.5 μg/mL. Capillary vessels were defined as round and hollow structures, localized in the periphery of muscle cells, and having a diameter of 12 μm maximum. Samples were analyzed with an OLYMPUS BX51 microscope and images from the total area of each histologic section were captured using an Evolution MP (Media Cybernetics, USA). The number of capillary vessels was determined in total muscle sections and the ratio of capillary vessels/mm^2^ and capillary vessels/muscle fibers were calculated using the Image Pro 6.3 software (Media Cybernetics, USA).

### Quantification of nerves

Groups of three mice were injected with *C. perfringens* JIR325 or with sterile PBS (negative controls) and sacrificed at 3 and 30 d post infection. Muscles were processed as described and embedded in paraffin. For each muscle, three non-consecutive sections of 4 μm were obtained and placed in positive Chargéd glass slides, then deparaffinized in xylene and hydrated. Antigen retrieval was performed using Proteinase K (Fermentas, Thermo Fisher Scientific) for 5 min at room temperature. For blockage steps, Dako Cytomation Biotin Blocking System (Dako, USA) as well as with Protein Block Serum-Free (Dako, Denmark) were used following the manufacturer’s instructions. Sections were incubated overnight with an Anti-200 kD Neurofilament Heavy Antibody-Neuronal Marker (Cat. No. ab8135; Abcam, USA) (diluted 1:1000) at 4°C in a wet chamber. Then, sections were washed with PBS and incubated with Polyclonal Goat Anti-Rabbit Immunoglobulins/Biotinylated (Cat. No. E0432; Dako, Denmark) (diluted 1:200) for one h at room temperature. After washing with PBS, sections were incubated with a Streptavidin Alexa Fluor 488 (Cat. No. S11223; Invitrogen) (diluted 1:200) for 30 min at room temperature. Finally, nuclear staining was performed with bis BENZIMIDE Hoechst (Cat. No. 33258; Sigma, USA) in a final concentration of 0.5 μg/mL. Images of the complete histologic sections were captured using an Evolution MP camera (Media Cybernetics, USA) in an OLYMPUS BX51 microscope. Structures between 50 and 8000 μm^2^ were considered and the number of nerves per mm^2^ and the number of axons per μm^2^ were determined with the Image Pro 6.3 software (Media Cybernetics, USA).

### Statistical analysis

Data were analyzed by the statistics softwares IBM^®^SPSS Statistics^®^ and GraphPad Prism version 5.00 for Windows. CK, PCR-RT and ELISAs were analyzed using Kruskall Wallis test and Dunn test as posthoc analysis. For analysis related to muscle regeneration process, area, muscle fibers, capillary vessels and nerves quantification, and the Mann-Whitney U test was used.

## Results and Discussion

### A sublethal inoculum of *C. perfringens* induces myonecrosis

*C. perfringens* is an anaerobic bacterium that induces gas gangrene, a devastating disease characterized by severe myonecrosis. Gas gangrene often occurs when vegetative or bacterial spores infect traumatic or chirurgical wounds and proliferate. We have previously shown that an intramuscular inoculum of 6×10^8^ CFU of *C. perfringens* induces myonecrosis, as evidenced by a rapid release of CK into the circulation (Monturiol-Gross, 2012). In this work, it was found that an inoculum of 1×10^6^ CFU of this bacterium also induces a significant increase in plasma CK 5 h after infection (p<0.01), although at 24h postinfection the plasma CK activity showed no significant difference compared to controls (Fig. 1A), indicating that the infection was controlled by the immune system and that the process of myonecrosis was limited in time. Histological analysis of the infected muscle showed areas of myonecrosis characterized by hyaline myofibrillar material and hypercontraction of myofibers since 5 h post infection (Fig. 1B). Bacterial aggregates were evident between the muscle fibers whereas the inflammatory infiltrate was distributed in a non-homogeneous way in the muscle, without accumulation of PMN inside the venules (Fig. 1B). Accordingly, it was previously reported that the inhibition of chemotaxis at the site of infection by *C. perfringens* depends on the size of the inoculum, and thus a sublethal inoculum does not effectively inhibit the inflammatory infiltrate into the infected muscle; moreover, the immune system is able to control the infection, inhibiting the establishment of the bacteria (O’Brien et al., 2007). Thus, our model of a sublethal inoculum of the bacteria allowed us to study the development of the muscle regenerative response.

**Figure 1.**
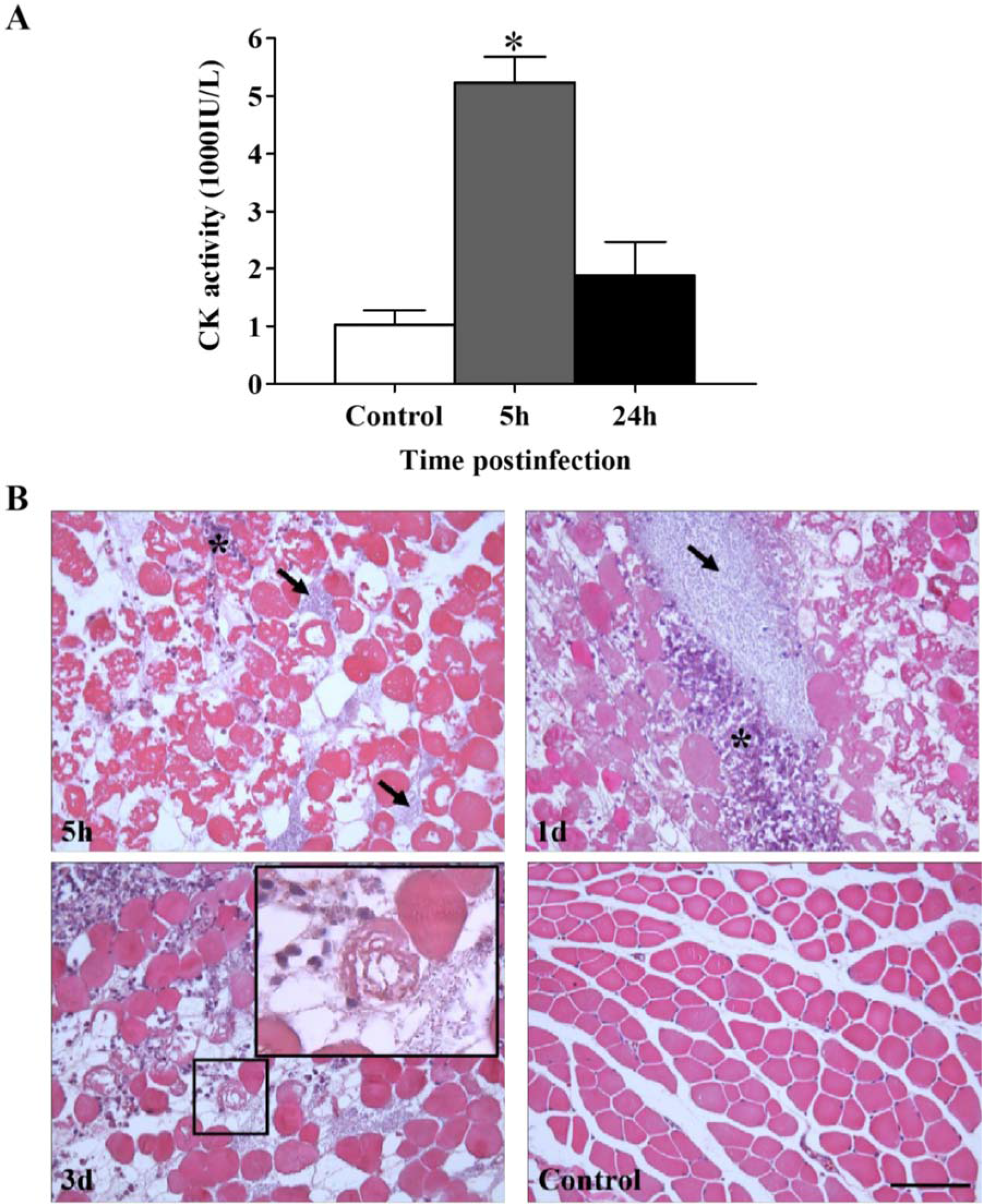
Myonecrosis induced by infection with a sublethal inoculum of *C. perfringens*. Groups of 6 CD-1 mice were injected intramuscularly with 1×10^6^ CFU of *C. perfringens*. **(A)** CK activity was determined in plasma 5 and 24 h after infection. Results show mean ± SE. *p<0.01 for samples that differ statistically from control. **(B)** Sections of the injected muscles were taken 5 h, 1 d and 3 d after infection, and stained with Hematoxylin-Eosin (HE). Abundant necrotic muscle cells are observed as early as 5 h after infection. Bacteria and inflammatory infiltrate are evident in the necrotic muscle (arrows and asterisks, respectively). Notice the presence of bacteria near a necrotic cell 3 d after infection (patch). Control muscle injected with PBS show a normal histological pattern. Bar scale, 100 μm.

### A sublethal inoculum of *C. perfringens* impairs the muscle regeneration process

To characterize the process of muscle regeneration, a histological analysis of the infected muscles was performed 7 d post infection. At this time, the presence of regenerative myofibers, characterized by central nuclei (Fig. 2), near or within a fibrous matrix, was observed. In addition, cellular detritus from necrotic fibers had not been removed despite the presence of an inflammatory infiltrate even at 14 and 30 d post infection (Fig. 2).

**Figure 2.**
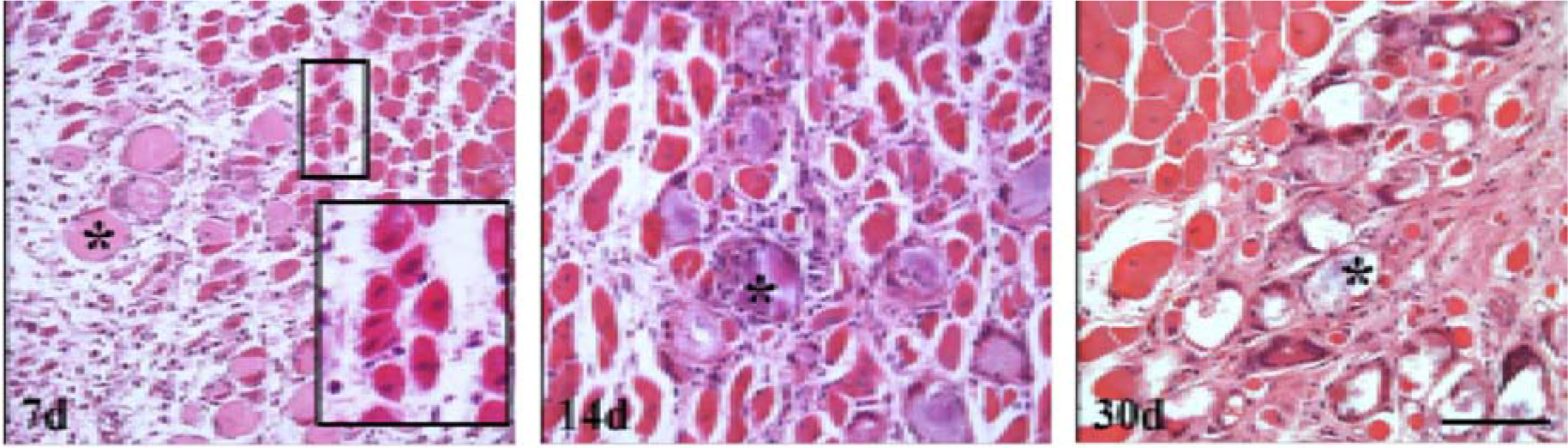
Alterations in muscle regeneration after an experimental infection with a sublethal inoculum
of *C. perfringens*. Groups of 3 CD-1 mice were injected intramuscularly with 1×10^6^ CFU of *C. perfringens* and muscle sections from samples collected at 7, 14, and 30 d were stained with HE. Patch at 7 d shows regenerative cells with central nuclei; asterisks indicate the presence of cellular debris embedded in a fibrous matrix at different times. Bar scale, 100 μm.

Due to the presence of regenerative cells, cell debris and fibrotic areas, regeneration and necrosis areas were quantified at 14 and 30 d after muscle injury (Fig 3A). At 24 h post infection the area of myonecrosis was 48.33 ± 6.5% of the total area of the muscle (initial lesion); at 14 d the percentage area corresponding to non-regenerated muscle was 22.12 ± 4.5%, while only 27.9 ± 5.3% of the muscle (Fig. 3A) was occupied by regenerating muscle fibers, hence evidencing a deficient muscular regeneration process.

**Figure 3.**
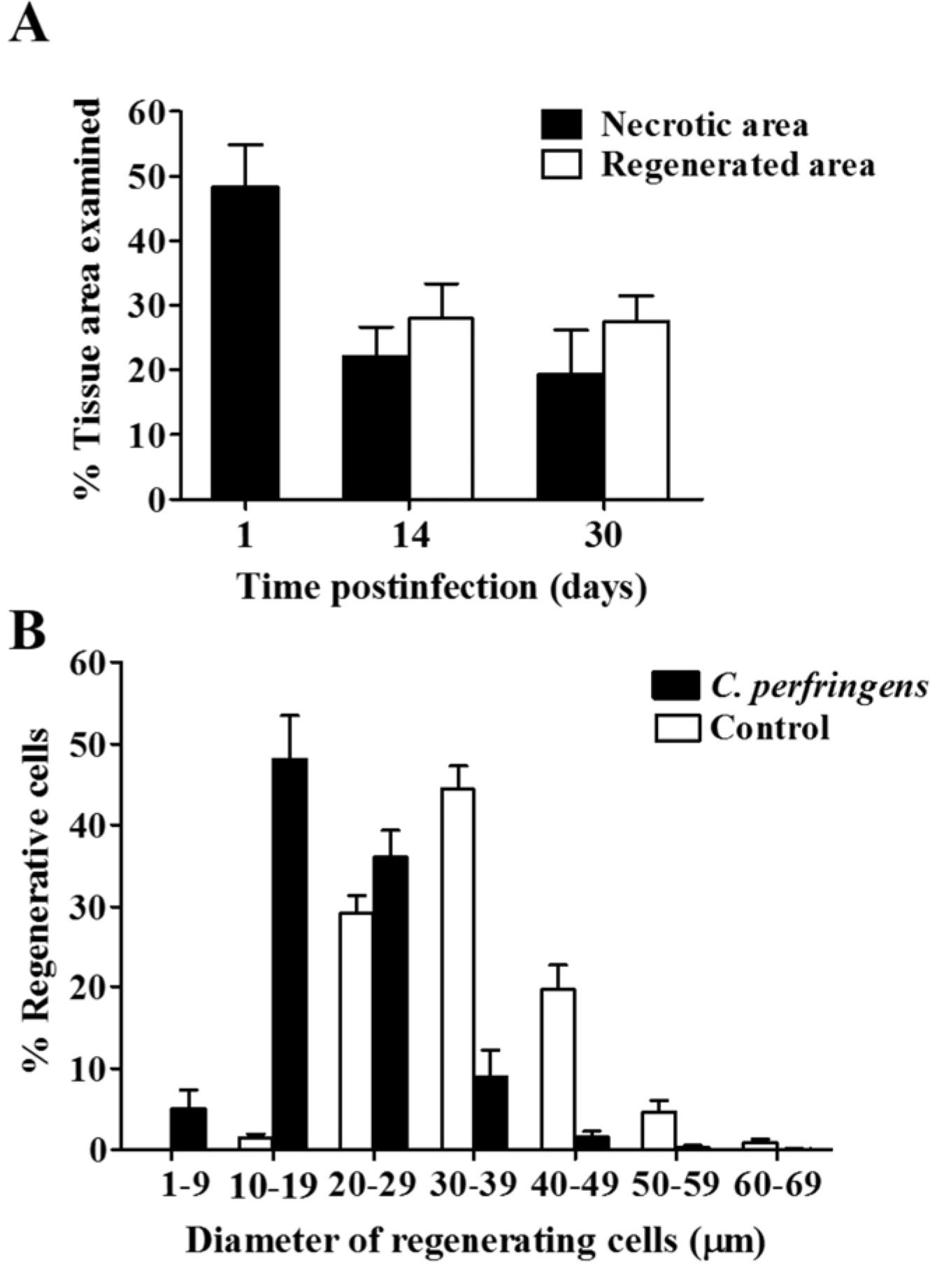
Muscle regeneration efficiency and regenerating fibers size after an experimental infection with a sublethal inoculum of *C. perfringens*. Groups of 3 CD-1 mice were injected intramuscularly with 1×10^6^ CFU of *C. perfringens*, and muscle sections from samples collected at 1, 14, and 30 d were stained with HE. **(A)** The extent of myonecrosis was determined one d post infection as the percentage of the examined area corresponding to necrotic fibers; the percentage of the necrotic area at 14 and 30 d corresponds to non-regenerated muscle including cellular debris and fibrotic zones, while the percentage of the regenerated area was determined 14 and 30 d post infection as the area encompassing regenerative fibers. **(B)** Quantification of the regenerative fibers according to their diameter 30 d post infection and comparison with controls injected with sterile PBS. Results show the means ± SE.

Although a portion of the damaged muscle was regenerated, when determining the size of the regenerative fibers it was observed that it differs significantly from the size of the fibers in control muscles at 30 d (Fig. 3B). While in the controls 44.5 ± 2.8% of the fibers had a diameter between 30-39 μm, in the muscles infected with *C. perfringens* 48.1 ± 5.4% of the fibers had a diameter between 10-19 μm. Moreover, 5.0 ± 2.4% of the regenerative cells in the infected muscles corresponded to fibers with a small diameter between 1-9 μm, indicating poor regeneration (Fig. 3B).

Another characteristic of poor regeneration is the replacement of muscle fibers by fibrous tissue. Specific stain for collagen fibers was made with Syrian Red. Red areas (indicative of collagen deposition) around small regenerative cells were evident 7 d post infection in the damaged muscle. These areas remained over time and showed greater intensity at 14 d and 30 d (Fig. 4A) after the muscle injury. When performing a quantitative analysis at 30 d, the control showed only 3.1 ± 0.5% of collagenous material in the total area, while in the muscles infected with a sublethal inoculum of *C. perfringens*, 23.5 ± 5.6% of the muscle corresponded to collagen deposition, which represents a significant difference between both groups (p <0.05) (Fig. 4B). Thus, the poor muscle regenerative outcome in this model correlates with an increased collagen deposition, underscoring the substitution of muscle fibers by a fibrotic matrix.

**Figure 4.**
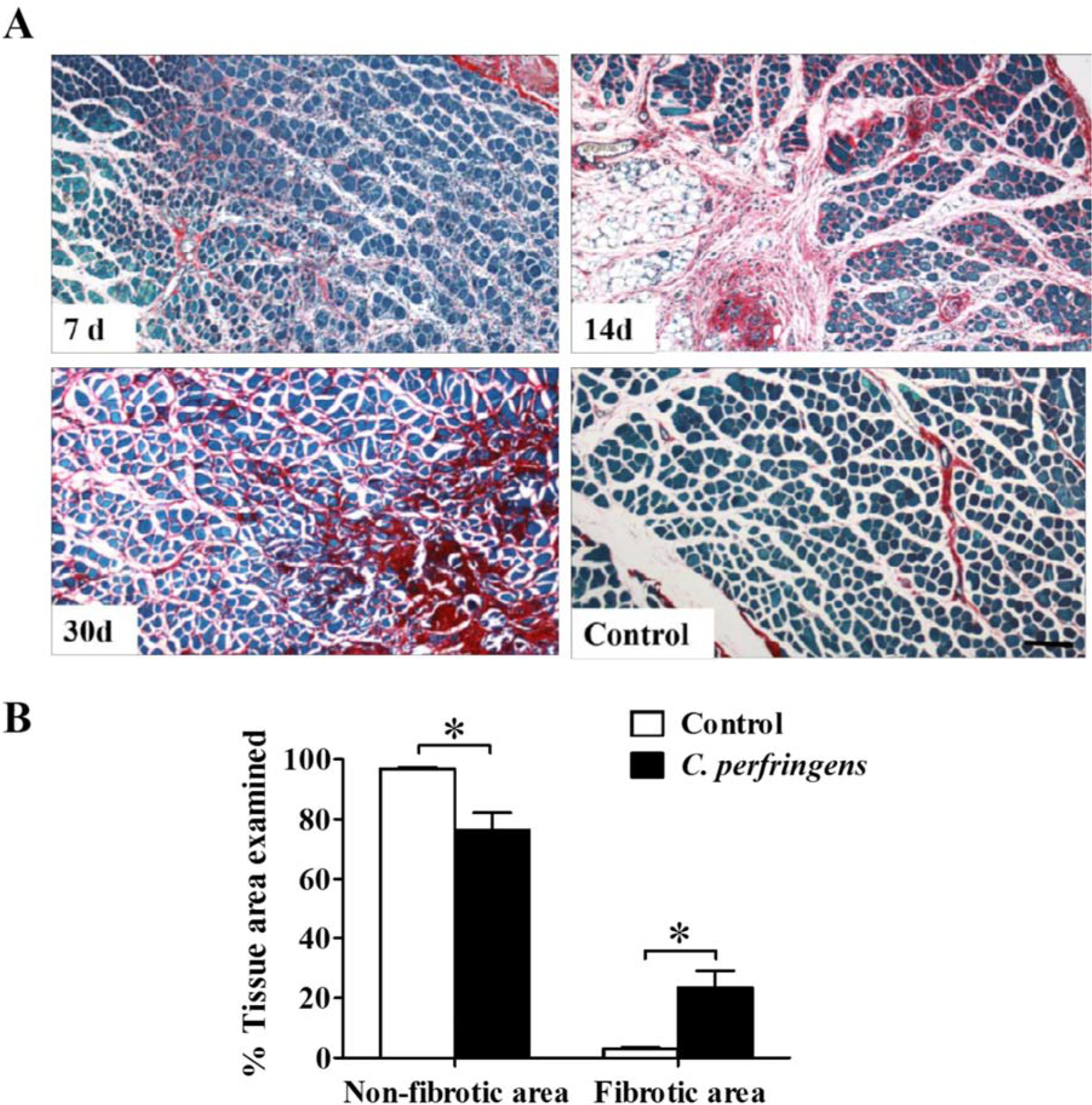
Collagen deposition in mice gastrocnemius after an experimental infection with a sublethal inoculum of *C. perfringens*. Groups of 3 CD-1mice were injected intramuscularly with 1×10^6^ CFU of *C. perfringens*. **(A)** Sections from muscle samples collected at 7, 14 and 30 d after injection were stained with Sirius Red and Fast Green; red areas correspond to collagen fibers, while green areas correspond to other proteins. Controls were injected with sterile PBS. Bar scale, 100 μm. **(B)** The fibrotic muscle was quantified at 30 d as the percentage of the examined area corresponding to collagen. Results show the means ± SE. *p<0.05 for samples with a statistically significant difference when compared with the control.

### A sublethal inoculum of *C. perfringens* alters capillary vessels and nerves in the infected muscle

To determine whether the infection with *C. perfringens* damages the vasculature, an immunostaining was performed with antibodies specific for the endothelial growth factor receptor Flk-1. At 6 h after the infection, a decrease in the number of capillary vessels was evident, and approximately 75% of the vessels were damaged 24 h after the infection with a sublethal inoculum of *C. perfringens* (Fig. 5A). The lack of capillary vessels was observed mainly in areas of myonecrosis 6 h and 1 d after the infection; in addition, the structures that showed a positive staining signal in these areas were smaller than those of the control samples, highlighting that they were probably non-functional capillary vessels or endothelial cell debris (Fig. 5A).

**Figure 5.**
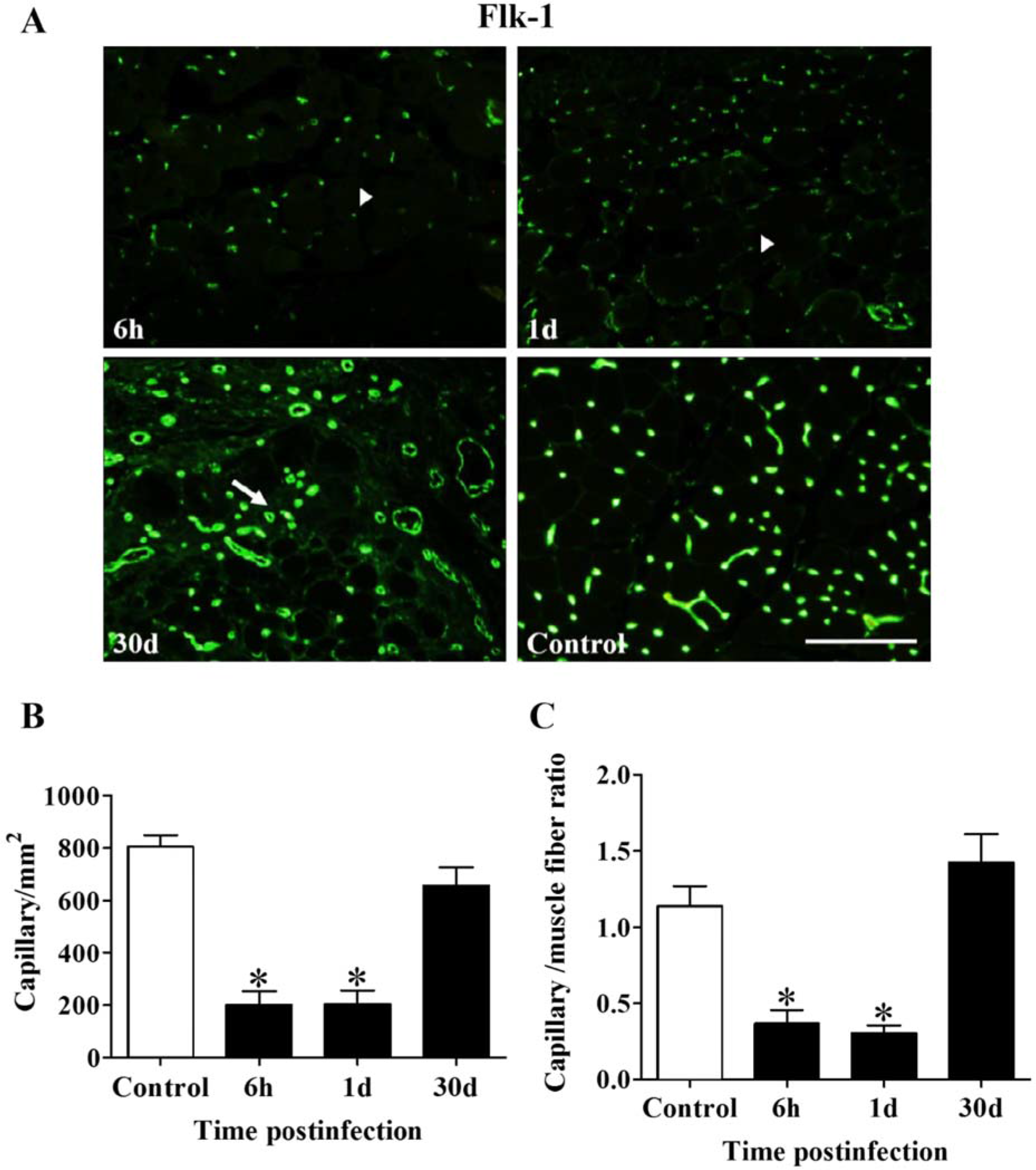
Capillary vessels in gastrocnemius after an experimental infection with a sublethal inoculum
of *C. perfringens*. Groups of 3 CD-1 mice were injected intramuscularly with 6×10^6^ CFU of *C. perfringens*. **(A)** Sections from muscle samples collected at 6 h, 1, and 30 d after injection were stained with anti-Flk-1 antibodies by immunofluorescence. Arrow heads indicate the lack of capillaries at 6 h and 1 d post infection, while arrows point the presence of capillaries in fibrotic areas at 30 d post infection. Bar scale, 100 μm. The number of capillaries per area **(B)** and per muscle fiber **(C)** were determined at different times after infection. Controls were injected with sterile PBS. Results show the means ± SE. *p<0.05 for samples with a statistically significant difference when compared with the control.

When the capillary vessels were quantified by area of tissue, the control showed an average of 805.3 — 43.8 capillaries per mm^2^, while in the muscles affected by *C. perfringens* the number decreased significantly to 201.3 ± 51.5 capillaries per mm^2^ at 6 h (p<0.05) and 203 ± 52.9 capillaries per mm^2^ at 24 h post infection (p<0.05) (Fig. 5B). Similar results were obtained when reporting the ratio of capillaries per muscle fiber, while the control showed an average of 1.14 ± 0.13 capillaries per muscle fiber, in the infected muscles the average significantly dropped to 0.37 ± 0.09 at 6 h (p<0.05) and to 0.30 ± 0.05 at 24 h post infection (p<0.05) (Fig. 5C).

Despite the significant decrease in the density of capillary vessels in early times after infection, at 30 d there were no significant differences in the number of capillaries per area or muscle tissue when compared to the controls (Figs. 5B and 5C), and they were evident even in the fibrotic tissue (Fig. 5A), suggesting a revascularization process. However, it is likely that the early disruption of the microvascular network in the infected muscle affects the process of regeneration, since key steps in muscle regeneration requiring an intact vascular supply occur within the first hours after myonecrosis. Muscle healing is critically affected by the ischemia associated with a deficient blood supply (Kotwal and Chien 2017). The most critical consequence of ischemia is a decrease in cellular energy supply (Kotwal and Chien 2017), as energy is required for every aspect of the wound-healing process such as protein synthesis, cell migration and proliferation, membrane transport, and growth factor production (Kotwal and Chien 2017). In these circumstances, the observed revascularization process may have occurred at a time when the muscle regenerative process had been already impaired.

Regarding the alteration of the nerves in the muscle, the antibody used against the neurofilament of the axons allowed their detection by immunofluorescence (Fig. 6A). A decreased number of nerves was observed 3 d after infection with *C. perfringens* (1.66 ± 0.27 nerves per mm^2^), when compared with the control (3.16 ± 0.11 nerves per mm^2^) (p<0.05) (Fig. 6B). In addition, a decreased number of axons within the nerves was observed (Fig. 6A); while in the control muscle the average was 17.03 ± 1.28 axons per 1000 μm^2^, in the affected muscles it decreased significantly to 6.72 ± 1.07 axons per 1000 μm^2^ (p<0.05) (Fig. 6C). Despite the damage observed 3 d post infection, when the samples were analyzed 30 d after muscle injury no significant differences were found in relation to the controls for the number of nerves per area nor for the number of axons per 1000 μm^2^ (Fig. 6C). Hence, a reinnervation process ensued in the muscle after the initial damage.

**Figure 6.**
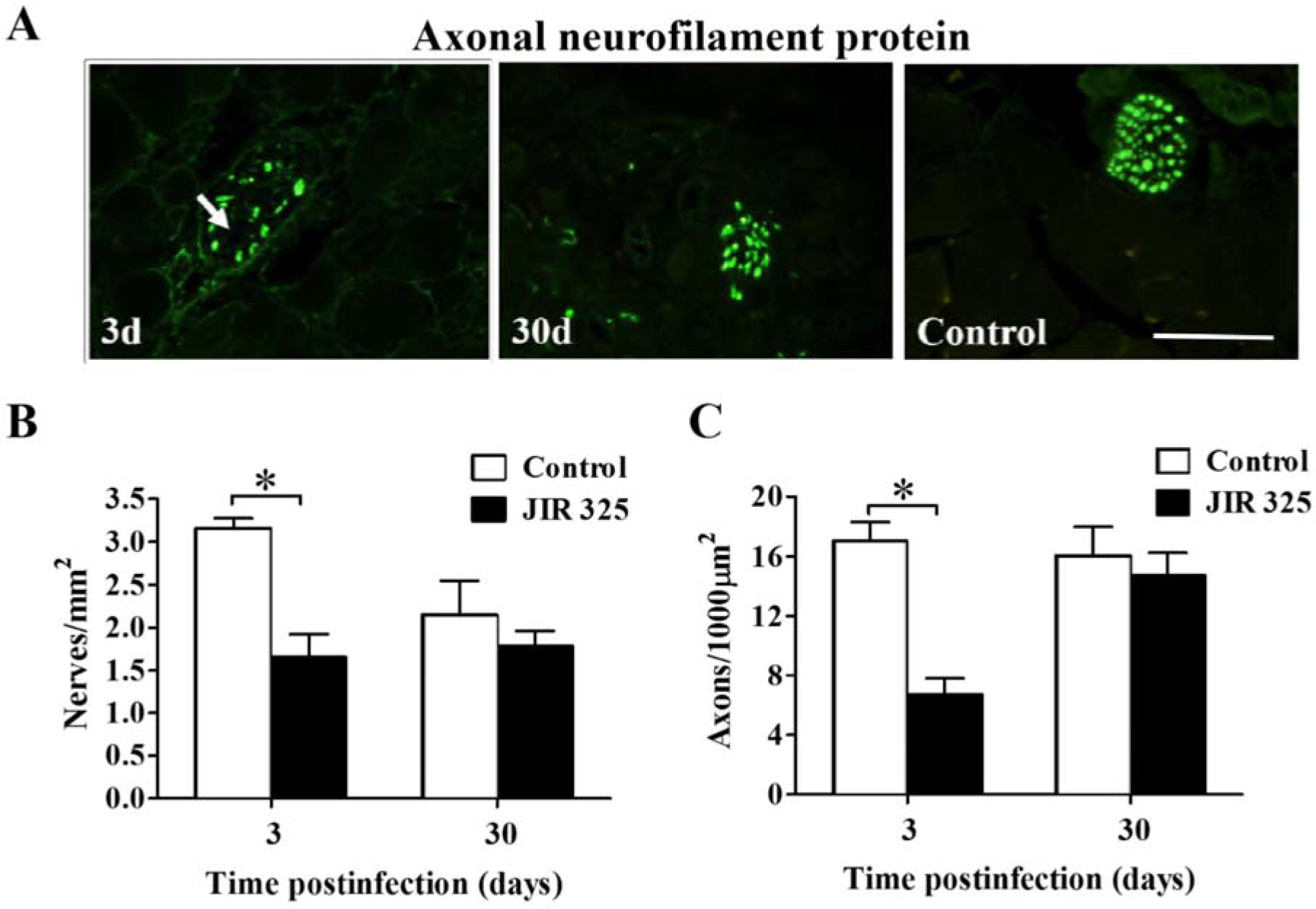
Innervation in mouse gastrocnemius after an experimental infection with a sublethal inoculum of *C. perfringens*. Groups of 3 CD-1 mice were injected intramuscularly with 6×10^6^ CFU of *C. perfringens*. **(A)** Sections from muscle samples collected at 3 and 30 d were stained with anti-heavy neurofilament protein antibodies by immunofluorescence. Arrow indicates nerve alterations 3 d post infection. Bar scale, 100 μm. The number of nerves per area **(B)** and the number of axons inside nerves **(C)** were determined at different times after infection. Controls were injected with sterile PBS. Results show the means ± SE. *p<0.05 for samples with a statistically significant difference when compared with the control.

### A sublethal inoculum of *C. perfringens* increases the expression of mediators of the inflammatory response and fibrosis in the infected muscle

To evaluate the immune response after infection with 110^6^ CFU of *C. perfringens*, expression of pro-inflammatory cytokines (IL1β, IL6, TNFα, IFNγ) and anti-inflammatory cytokines (IL4, IL10, IL13) was determined by RT-PCR and ELISAs (Fig. 7). Furthermore, the expression of PMN (MIP2 and KC), and macrophage (MCP-1) chemoattractant cytokines was also analyzed (Fig. 8).

**Figure 7.**
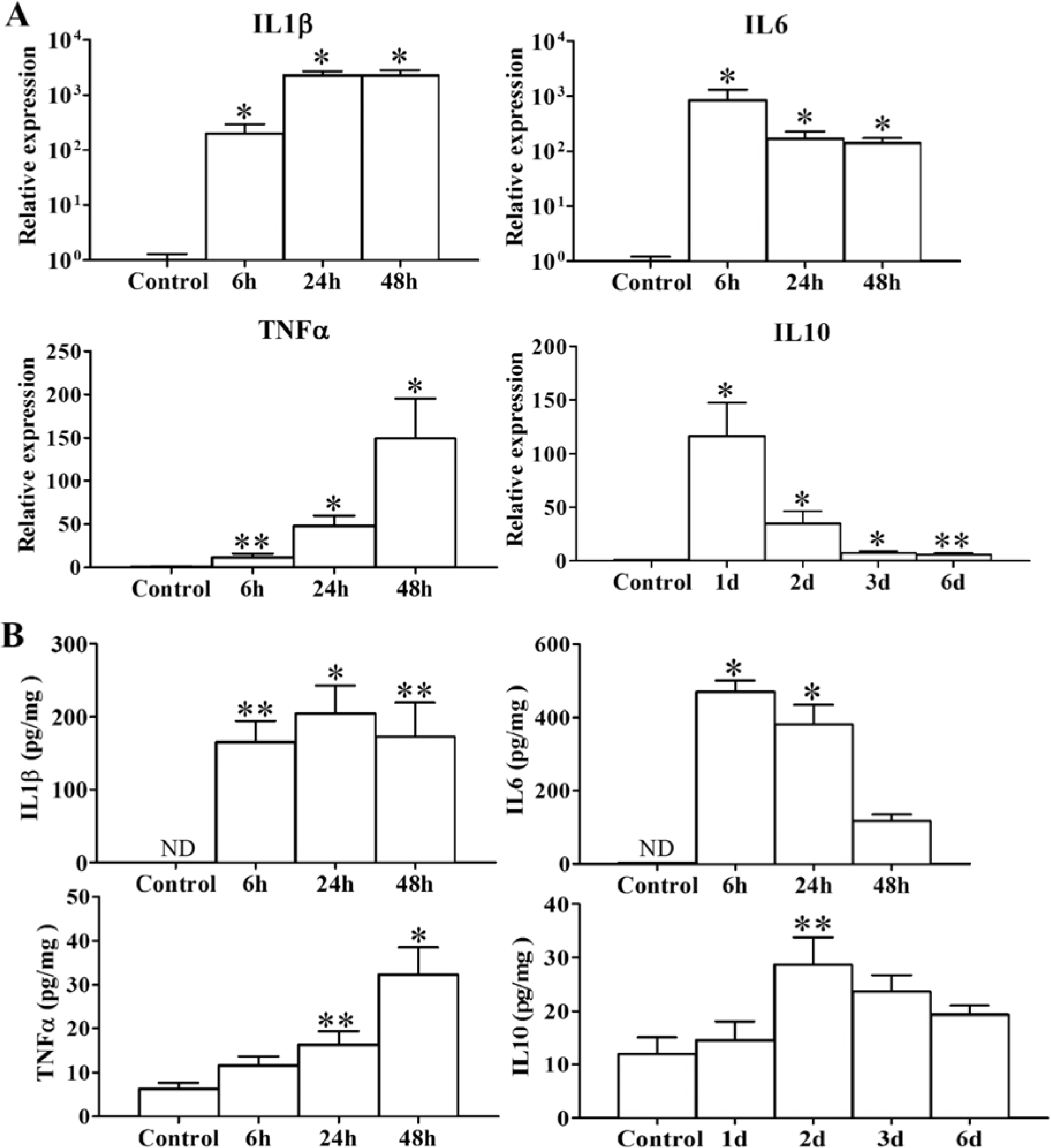
Cytokines expression in mouse gastrocnemius after an experimental infection with a sublethal inoculum of *C. perfringens*. Groups of 4 CD-1 mice were injected intramuscularly with 1×10^6^ CFU of *C. perfringens* and the expression of cytokines in the infected muscles was measured by RT-PCR **(A)** and by ELISAs **(B)** at different times. The results obtained with three normalization genes (GADPH, RPL13A, RNSP1) were incorporated in **(A)**, based on 2^−ΔΔCt^ calculation. Controls were injected with sterile PBS. Results show the mean ± SE. *<0.01; **p<0.05 for samples with a statistically significant difference when compared with controls.

**Figure 8.**
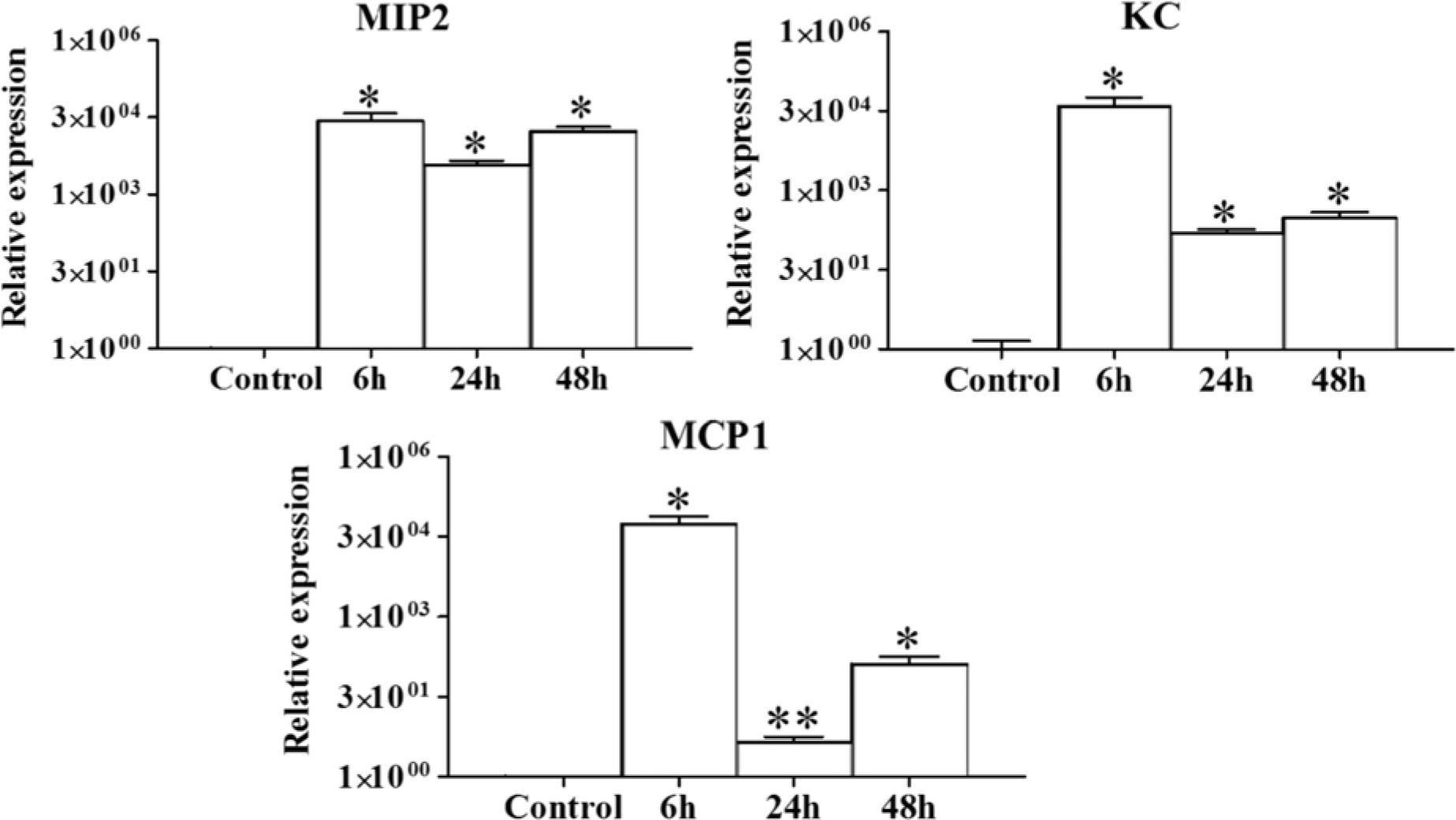
Chemoattractants expression in mouse gastrocnemius after an experimental infection with a sublethal inoculum of *C. perfringens*. Groups of 4 CD-1 mice were injected intramuscularly with 1×10^6^ CFU of *C. perfringens* and the expression of chemoattractans in the infected muscles was measured by RT-PCR at different times. The results obtained with three normalization genes (GADPH, RPL13A, RNSP1) were incorporated, based on 2^-△△Ct^ calculation. Controls were injected with sterile PBS. Results show the mean ± SE. *<0.01; **p<0.05 for samples with a statistically significant difference when compared with controls.

A significantly increased expression of mRNA for IL1β and IL6 was observed 6 h post infection when compared to controls, which lasted up to at least 48 h (p<0.01) (Fig. 7A). The expression of TNFα also showed a significant increase from 6 h on (p<0.05) and remained elevated for at least 48 h post infection (p<0.01) (Fig. 7A). Furthermore, a significant increase in the expression of IL10 was observed in comparison to controls from 1d (p<0.01) up to 6 d (p<0.05), reaching a maximum peak 24 h after infection with the bacteria (p<0.01) (Fig. 7A). When immunoassays were carried out to quantify various cytokines in the muscle, confirmatory results were obtained since IL1β remained high for at least 48 h post infection, IL6 increased significantly from 6 to 48 h post infection (p<0.01), reaching maximum level at 6 h, while the TNFα reached a maximum level 48 h post infection (p<0.01). In contrast, IFNγ showed only a significant increase at transcriptional level 6 h post infection (p<0.01), but not at the protein level (data not shown).

There was not a significant difference in the expression of IL4 between infected mice and controls, whereas for IL13, there was a significant difference only at 6 d post infection (p<0.05) (data not shown). On the other hand, IL10 showed a significant increase 2 d after infection (p <0.05) and the amount of protein remained slightly higher than controls until 6 d (Fig.7B).

The chemokines MIP2 and KC showed a significant increase in expression in comparison with controls from 6 h to 48 h post infection with a sublethal inoculum of *C. perfringens* (p<0.01) (Fig. 8). KC highest expression occurred 6 h post infection (maximum peak) and remained high even at 48 h. MCP-1 highest expression occurred 6 h post infection, although it decreased 24 h post infection and increased again at 48 h; its expression was significantly higher than that observed in the controls in all the evaluated times (Fig. 8).

Because the TGFβ1 has been associated with fibrosis in the process of muscle regeneration, its gene expression and protein concentration were analyzed. A bimodal behavior was shown for TGF β1 both at the transcriptional and protein levels. When the relative mRNA expression was analyzed, a significant increase was observed one day after infection (p<0.05) in relation to the control, its expression decreased at 2 d but increased again at 3 d (p <0.01), remaining elevated at least until 6 d post infection (p <0.05) (Fig. 9A).

**Figure 9.**
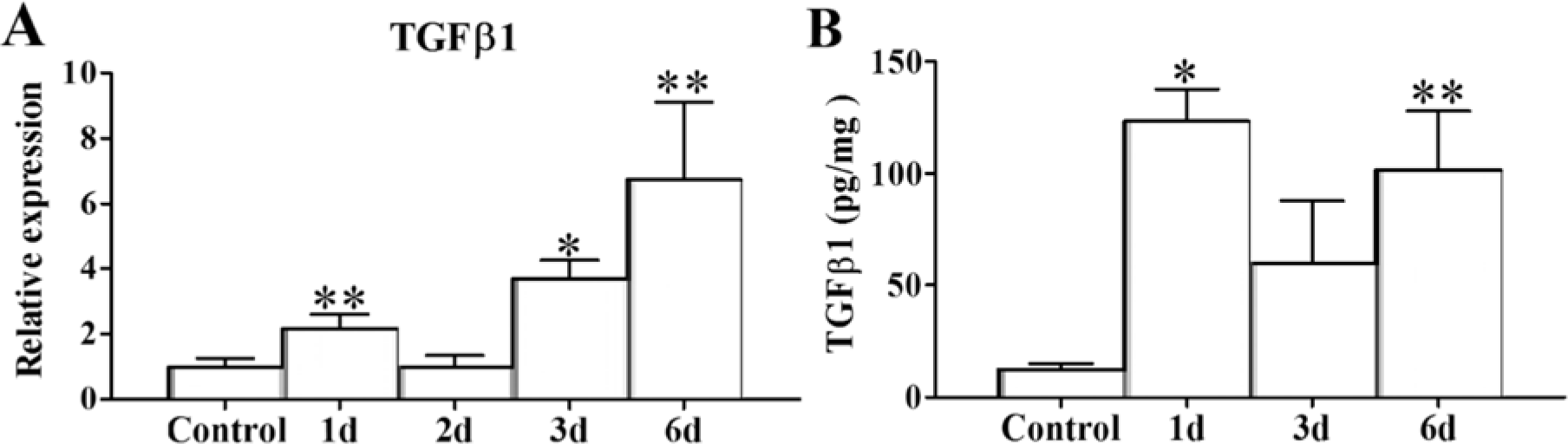
TGFB1 expression in mouse gastrocnemius after an experimental infection with a sublethal inoculum of *C. perfringens*. Groups of 4-6 CD-1 mice were injected intramuscularly with 6×10^6^ CFU of *C. perfringens* and the expression of TGFpi in the infected muscles was measured by RT-PCR **(A)** and by ELISA **(B)**, at different times. Controls were injected with sterile PBS. Results show the means ± SE. *p<0.01; **p<0.05 for samples with a statistically significant difference when compared with the control.

At the protein level, a significantly higher amount of TGFβ1 was detected in the infected muscle when compared to the control 1 d after infection (p<0.01), and although there was a decrease at 3 d, the protein concentration remained elevated 6 d post infection in the infected muscles, as compared to the controls (p<0.05) (Fig. 9B). Thus the increase in TGFβ1 correlates with the poor regeneration found after the infection with a sublethal dose of *C. perfringens.* This fits with the known role of this mediator which favors collagen deposition, i.e. fibrosis, and inhibits myogenic cell differentiation.

### A sublethal inoculum of *C. perfringens* alters the influx of PMN and macrophages to the infected muscle

PMN are the first cells that reach the muscle after a myonecrosis, and are followed by M1 and M2 macrophages before the resolution of muscle damage (Tidball, 2017). In injuries induced by a lethal inoculum of *C. perfringens*, the absence of inflammatory cells in the infected muscle and the presence of PMN attached to the endothelium due to the overexpression of adhesion molecules has been reported (Stevens and Bryant, 2002). However, when using a sublethal inoculum of this bacterium, an inflammatory infiltrate was observed in the muscle, with the presence of PMN, both aggregated and dispersed in the necrotic muscle (Fig. 10A). The presence of PMN was evident 5 h after infection (Fig. 10A), when they reached a number of 505.2 ± 38 cells per mm^2^ (Fig. 10B). At 24 h their number was 548.4 ± 56.40 cells per mm^2^, at 3 d 342.2 ± 36.57 cells per mm^2^, and at 5 d after infection numbers dropped to 81.68 ± 9.87 cells per mm^2^ (Figs. 10A and 10B). These observations agree with the described pattern of early neutrophil influx as the first wave of inflammatory cells in injured tissues.

**Figure 10.**
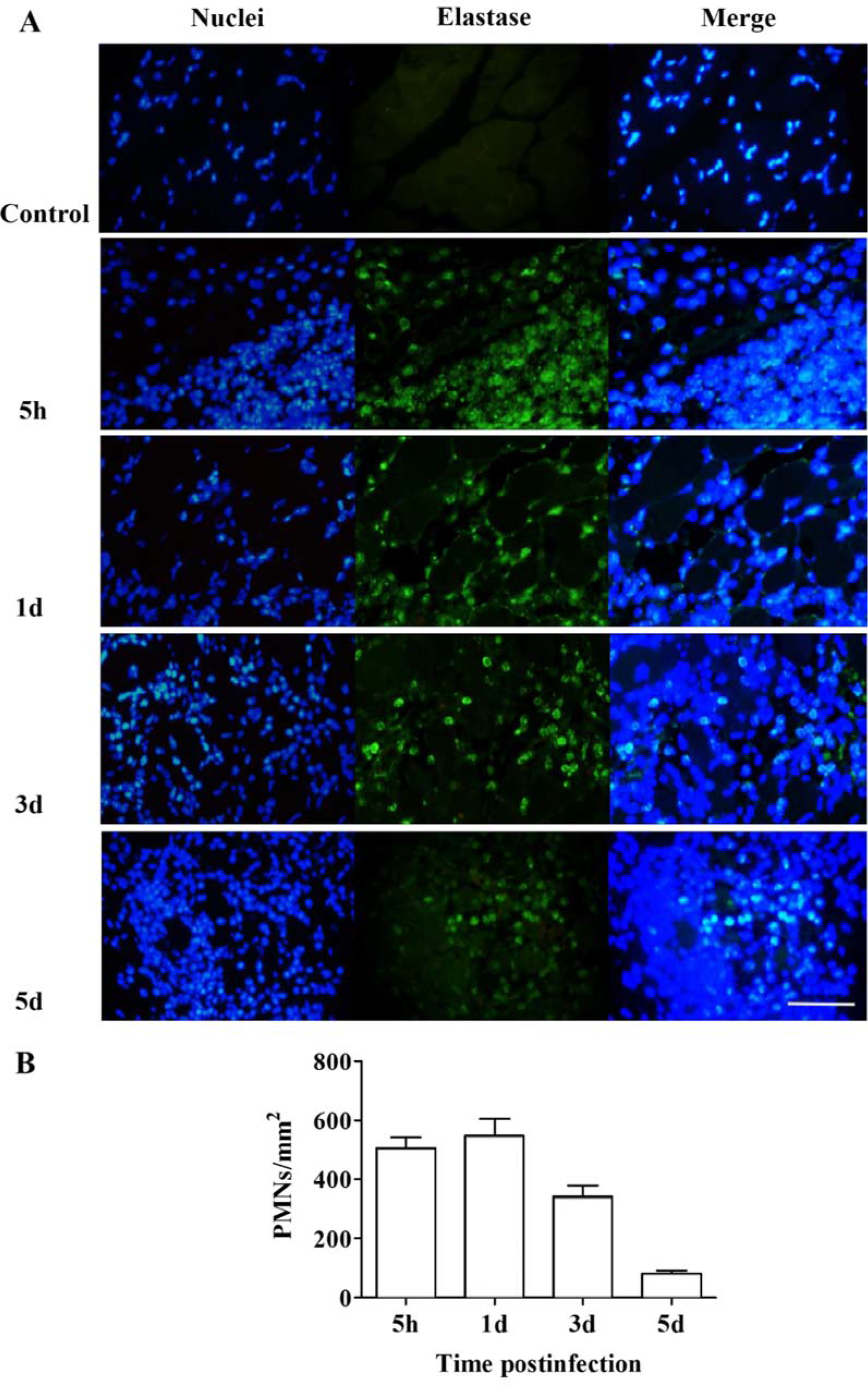
Recruitment of PMN at the site of injury after an experimental infection with a sublethal inoculum of *C. perfringens*. Groups of 3 CD-1 mice were injected intramuscularly with 1×10^6^ CFU of *C. perfringens* and the presence of PMN was determined using anti-elastase antibodies at the indicated times **(A)**. For each time, the number of cells corresponding to PMN was determined **(B)**. Bar scale, 50 μm.

M1 macrophages immunostained with anti-iNOS were poorly detected during the study period. Its density was 106.3 ± 23.18 M1 cells per mm^2^ 24 h post infection, reached maximum at 2-3 d post infection (142.60 ± 18.21 cells per mm^2^) (Figs.11A and 11B) and decline further until 5 d post infection (25.43 ± 8.51 cells per mm^2^) (Fig. 11B). M2 macrophages immunostained with anti-arginase antibodies were detected 24 h post infection and reached a maximum density 7 d post infection (616.2 ± 179.4 cells per mm^2^) (Figs. 11A and 11B), although they were detected even 14 d post infection (41.82 ± 18.06 cells per mm^2^) (Fig. 11B).

**Figure 11.**
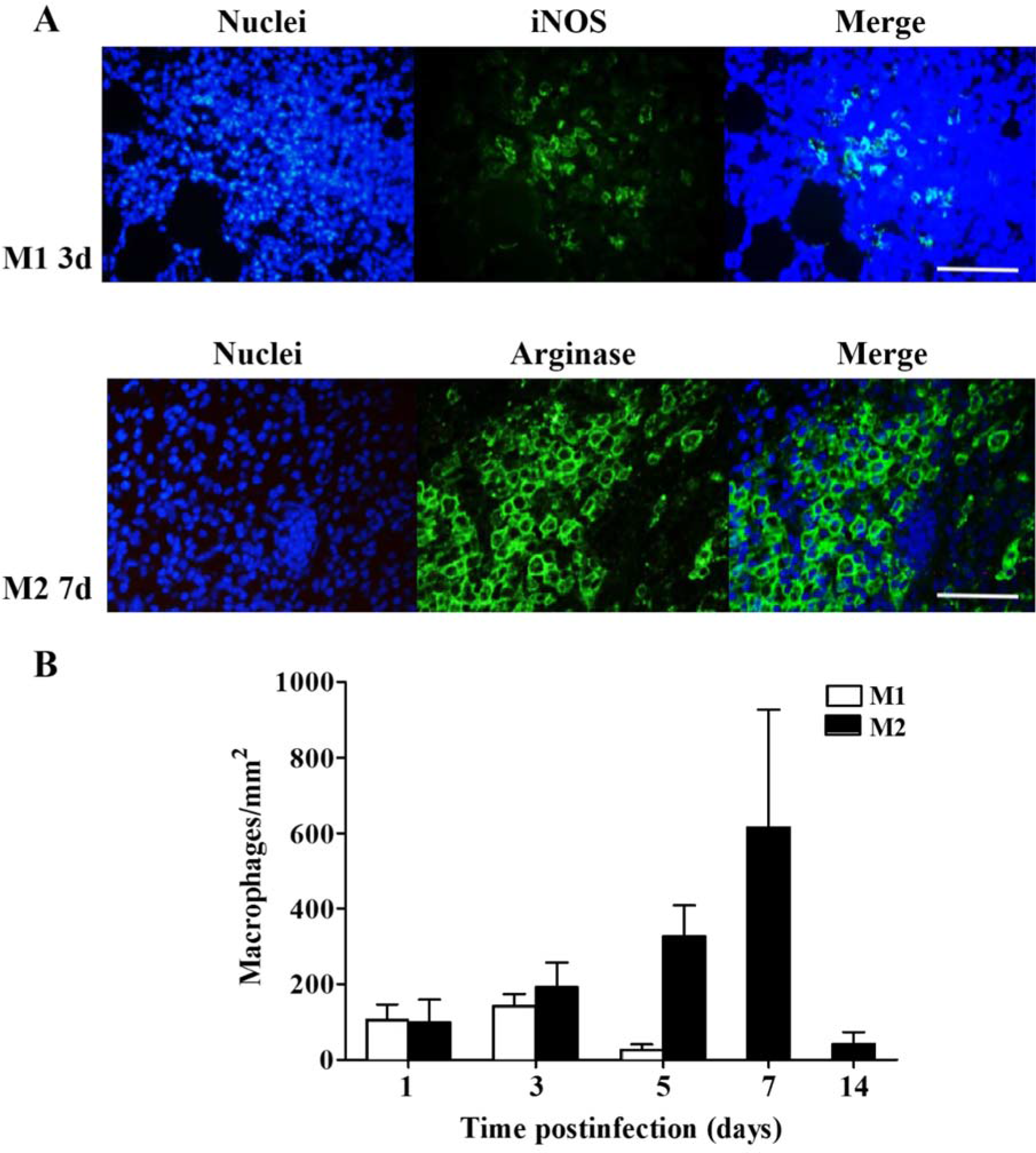
Recruitment of M1 and M2 macrophages at the site of the injury after an experimental infection with a sublethal inoculum of *C. perfringens*. Groups of 3 CD-1 mice were injected intramuscularly with 1×10^6^ CFU of *C. perfringens* and the presence of M1 and M2 macrophages was determined using anti-iNOS and anti-arginase antibodies, respectively, at the indicated times. The highest intensity observed was related to the maximum number of cells in the muscle **(A)**. For each time, the number of cells corresponding to M1 and M1 macrophages was determined **(B)**. Bar scale, 50 μm.

Little influx of M1 macrophages (iNOS^+^) in the muscle of the infected animals was evident (Figs. 11A and B), which suggests alterations in this population of inflammatory cells after infection with a sublethal inoculum of *C. perfringens.* This could be due to the observed absence of IFN-γ. Accordingly, in experiments in which IFNγ signaling has been blocked in injured muscles, there is a reduction in the expression in macrophages of transcripts that indicate the activation of the M1 phenotype, such as iNOS (Cheng et al., 2008). The classical inflammation response after a tissue injured occurs within the following 5 d (Novak and Koh, 2013). Normal remodeling in muscle is very dependent on the timing of the M1 and M2 macrophages response (Chazaud, 2016; Xiao et al., 2016). The transition to M2 macrophages is critical to muscle regeneration, and prolonged inflammation results in fibrosis (Chazaud, 2016; Xiao et al., 2016).

IL10, IL4 and IL13 have defined roles in the activation of M2 macrophages, which constitute a complex population comprising several subpopulations with functional and molecular specializations (Mantovani et al., 2004; Tidball and Villalta, 2010). M2a macrophages are activated by IL4 and IL13, secrete IL10, express arginase 1, and promote wound healing and muscle regeneration, while M2b are activated by immune complexes or Toll-like receptors, and release anti-inflammatory cytokines associated with the Th2 response (Tidball and Villalta, 2010; Rigamonti, 2013). On the other hand, M2c macrophages are activated by IL10, release cytokines that deactivate the M1 phenotype, promote the proliferation of nonmyeloid cells, and influence the deposition of the extracellular matrix (Tidball and Villalta, 2010; Tidball, 2017). Hence, the dynamics of cytokine synthesis in the affected muscle greatly determines the appearance and action of various populations of macrophages in a complex tissue landscape which, in turn, greatly determines the outcome of the regenerative process. Macrophages play a central role in the natural wound healing (Juban and Chazaud, 2017). They direct T-cell activation, promoting stem cell and progenitor cell migration, activating angiogenic responses, and guiding extracellular matrix remodeling (Castiglioni et al., 2015). They phagocytose cellular debris generated during tissue remodeling, recycling important molecular components to be reused (Juban and Chazaud, 2017). The phenotype of the macrophages can have profound influences on the progression of disease or injury (Juban and Chazaud, 2017). The same macrophage can switch between pro-inflammatory and pro-healing states depending on the surrounding environmental cues, and the switch from the M1 to the M2 phenotype occurs in stages in response to upregulation of IL-4 and IL-13 (Juban and Chazaud, 2017).

Recruited monocytes fail to differentiate adequately in tissue remodeling, in muscular dystrophy (Villalta et al. 2009), insulin resistance (Olefsky and Glass 2010), and advanced age (Mahbub et al. 2012) leading to unsuccessful remodeling and tissue repair (Carvalho et al. 2013; Villalta et al. 2011a, b). Similar to muscular dystrophies, muscle loss due to aging is likely caused by a prolonged inflammatory response, characterized by a higher expression of IL-1β. If the M1 response lasts too long, the new tissue is highly fibrotic, leading to a decreased function. In myonecrosis induced by a subletal inoculum of *C. perfringens* altered muscle regeneration was observed in association to both a delayed recruitment of macrophages at the site of infection and an altered production of cytokines.

After infection with a sublethal inoculum of *C. perfringens*, a significant increase in the gene expression level of IL10 in the muscle was observed from 1 d (maximum peak) to 6 d after infection, and at the protein level since 2 d (maximum peak) (Figs. 7A and B). Although no significant changes were detected in the expression of IL4, for IL13 an increase in gene expression level was observed after 6 d (data not shown). The influx of both M1 and M2 macrophages into the infected muscle was observed since the first day post infection. The presence of M1 macrophages in the infected muscle was scarce, having its maximum peak by 3 d whereas M2 mecrophages were more abundant than M1 in the infected muscle, having its maximum peak by 7 d and remaining high even after 14 d (Fig. 11B). The altered arrival of M1 and M2 macrophages into the infected muscle after infection in our model is likely to alter the muscle regeneration process.

When analyzing the arrival and permanence of the different cell populations using a sublethal dose of *C. perfringens* vis-à-vis the typical inflammatory response reported in the literature (Fig. 12), there is a prolongation of the inflammatory response in the *C. perfringens* model. The PMN remain in the infected muscle up to 5 d, which is complemented with a high expression of proinflammatory cytokines for up to at least 2 d. In addition, there is a limited arrival of M1 macrophages which remain within the infected muscle for longer time span than expected. It is proposed that given the absence of IFNγ in the acute phase of infection, the number and activity of M1 macrophages were affected. One possible explanation for this could be an altered function of PMN due to the direct effect of *C. perfringens* toxins in these cells. It has been reported that perfringolysin O (PFO) induces cytotoxicity in PMN (Stevens and Bryant, 2002) and, *C. perfringens* phospholipase C (CpPLC) interferes in the replacement of mature PMN in the peripheral circulation, inhibiting their maturation (Takehara et al., 2016). Additionally, the vascular damage caused by *C. perfringens* infection could also contribute to prevente the influx of cells of the inflammatory response. Thus, the direct and indirect effects of the infection with *C. perfringens* on inflammatory cells results in evident consequences in the regenerative response.

**Figure 12.**
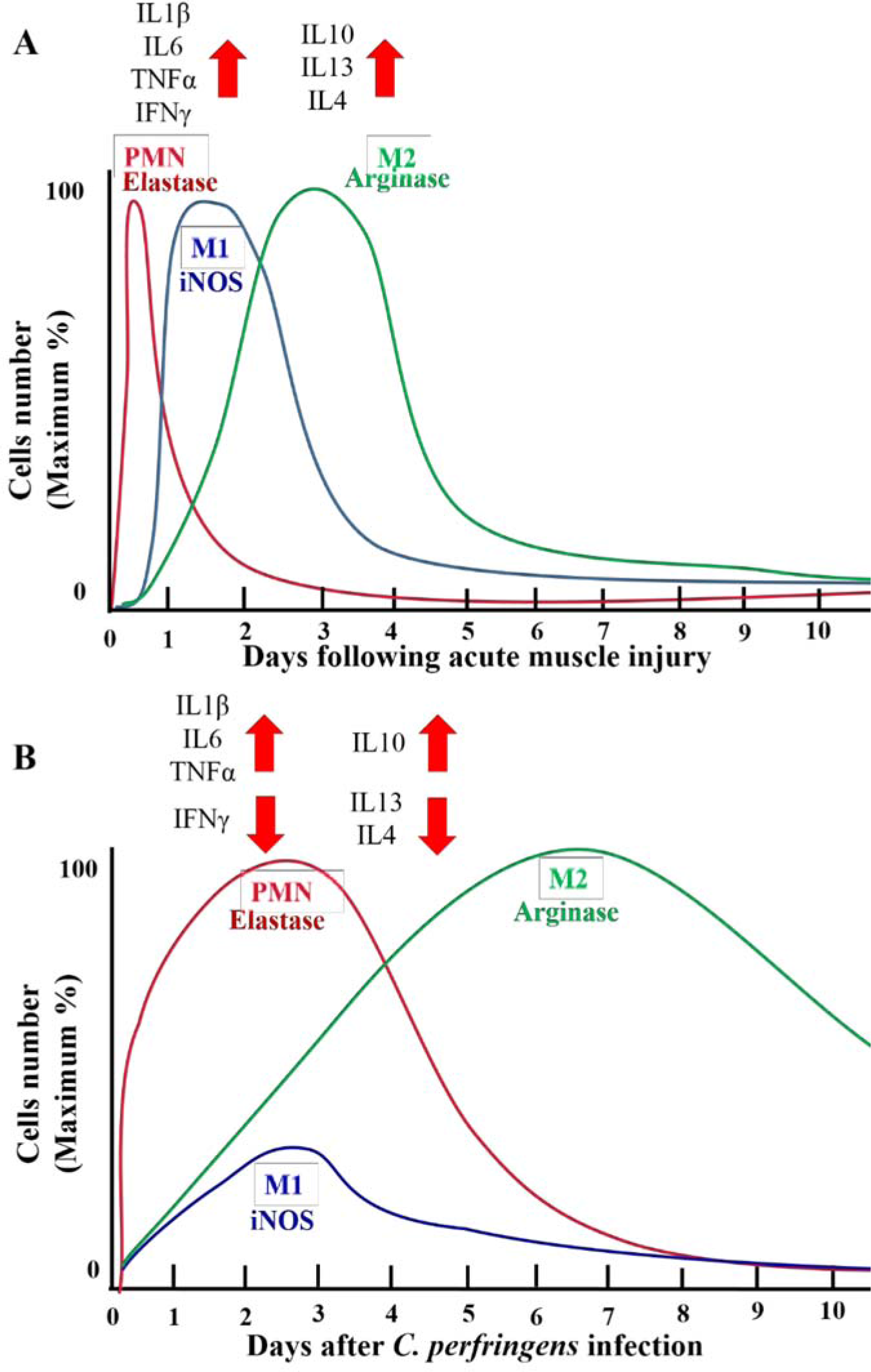
Time course of changes in cytokines expression and intramuscular myeloid cell populations after acute muscle injury, compared with those after an experimental infection with a sublethal inoculum of *C. perfringens*. (A) Expected cytokines expression and changes of myeloid cell populations after acute muscle injury (Tidball and Villalta, 2010; Tidball, 2017). (B) Cytokines expression and changes of myeloid cell populations after an intramuscular infection with a sublethal inoculum of *C. perfringens*. PMN, neutrophils; M1, M1 macrophages; M2, M2 macrophages.

After a muscle injury, the regeneration process is mediated by a specific type of stem cells, the SC, but also involves the interaction of these myogenic cells with other resident cells, inflammatory cells, blood vessels, nerves and the extracellular matrix (Ciciliot and Schiaffino, 2010; Hernández et al., 2011). There are at least three basic requirements for the process of muscle regeneration to occur: a) an adequate blood flow in the regenerative muscle; b) innervation of regenerative cells; and c) permanence of the basal lamina around the necrotic muscle fibers, which serves as a scaffold and substrate for regeneration (Gutiérrez et al., 2018). Impairment of any of these contributory factors results in a defective muscle regeneration.

The importance of blood flow lies, on one hand, in the inflow of inflammatory infiltrate to the site of the lesion and, in addition, to the provision of oxygen and nutrients and ATP to the regenerating muscle. When vascular density was analyzed after an infection with a sublethal inoculum of *C. perfringens*, it was observed that the microvasculature is affected in the first hours after infection, a time that coincides with the arrival of PMN, the first inflammatory cells present in the infected muscle. Although the presence of the inflammatory infiltrate is evident, the vascular damage and the direct effect of CpPLC and PFO on cells of the immune system can affect the migration of inflammatory cells to the area of damage, which results in the deficient removal of necrotic debris. Thus, the combination of a direct inhibitory action of *C. perfringens* toxins, added to the disruption of the microvascular network, is likely to affect the timely arrival of inflammatory cells. It has been reported that disturbances that affect the removal of dead cells delay the process of muscle regeneration (Tidball, 2017; Zhao et al., 2016). This may be due to the fact that the persistence of cellular debris becomes a physical obstacle for muscle regeneration.

Another consequence of the damage to the vascular system in the regenerative process is the limitation in the available oxygen, which is critical for muscle healing. Hypoxia can also promote the proliferation of bacteria and the further development of gas gangrene, hence generating a vicious cycle (Flores-Díaz and Alape Girón, 2003). In studies carried out with snake venoms, the alteration of the microvasculature affects the regenerative process, favoring the replacement of muscle by fibrotic muscle; moreover, regenerative fibers have small diameters (Gutiérrez et al., 1984; Arce et al., 1991; Hernández et al., 2011). Although our findings showed that the capillary density is restored at 30 d, it is possible that the severe blood vessel damage induced by *C. perfringens* in the first hours after infection (Fig. 5) could be one of the causes behind the impaired regenerative process.

Innervation is another requirement for a successfull muscle regeneration process. The proliferation and fusion of myogenic cells occur in muscles that regenerate in the absence of nerves and in those in which the nerves are intact (Slater and Schiaffino, 2008). However, although the initial events of the muscle regeneration process can occur in the absence of innervation, the latter is required for the growth and recovery of muscle function (Kalhovde et al., 2005; Slater and Schiaffino, 2008). In the absence of innervation, regenerating cells do not reach their maturity. In the model, although the density and structure of the nerves were affected with a sublethal inoculum of *C. perfringens* 3 d after infection, innervation was recovered at 30 d (Fig. 6B). Consequently, innervation does not appear to be a cause of poor regeneration after muscle damage generated by *C. perfringens.* However, the question remains as to whether *C. perfringens* toxins induce damage at the synaptic level and neuromuscular transmission, as has been reported for the lethal toxin of *Clostridium sordelii* (Barbier et al., 2004). For a successful muscle regeneration process, scaffolding is required to maintain the position of the muscle fibers.

The processes of muscle necrosis and regeneration involve a complex turnover of the extracellular matrix, which is determinant for an effective regenerative response. The process of matrix deposition after myonecrosis, although initially beneficial, affects the regenerative process if it continues without control, resulting in the permanent accumulation of collagen around the myofibers, which may even lead to muscle replacement by fibrous tissue (Serrano and Muñoz-Cánoves, 2010). Fibroblasts contribute to the formation of fibrous muscle by the production and accumulation of components of the extracellular matrix, such as hyaluronic acid, fibronectin, proteoglycans and interstitial collagens (Serrano and Muñoz-Cánoves, 2010). When the histological sections of animals infected with a sublethal inoculum of *C. perfringens* were analyzed, it was observed that at 30 d, almost 25% of the muscle area corresponded to collagen accumulated in areas where the regenerative process was deficient (Fig. 4). This may be associated with the overexpression of some mitogenic chemokines and cytokines, produced by macrophages and PMN, which are also involved in the mechanism of fibrogenesis. It has been reported that MCP-1 is a profibrotic mediator, whose neutralization reduces the extent of fibrosis (Deshmane et al., 2009; Wynn, 2008).

Experimental models of fibrosis have documented potent antifibrotic properties for cytokines associated with the Th1 response such as IFNγ (Wynn, 2008); thus, the absence of this cytokine in our study model could be associated with overproduction of extracellular matrix. On the other hand, IL10 and TGFβ activate a subpopulation of M2 macrophages that promotes the deposition of extracellular matrix and fibrosis in different pathogenic conditions (Serrano and Muñoz-Cánoves, 2010; Tidball, 2017). TGFβ has been associated with the development of fibrosis in a number of diseases as it is one of the main activators of macrophages and fibroblasts; of the three isotypes of TGFβ in mammals, muscle fibrosis is mainly attributed to the TGFβ1 isoform (Yoshimura et al.2010; Wynn, 2008). In this work an increase of TGFβ1 was evidenced both in early and late time intervals (Fig. 9), and hence may be related to the collagen accumulation evidenced since 7 d.

IFNγ has a relevant function in muscle regeneration, since this molecule directly regulates myogenic cells in their differentiation process (Cheng et al., 2008; Tidball et al., 2017). Therefore, the alterations in the regenerative response observed in this study could also be a consequence of the absence of IFNγ. In general, infection with a sublethal inoculum of *C. perfringens* stimulates the expression of chemoattractants and cytokines that stimulate the arrival of different inflammatory cells at the site of the lesion. However, it has been previously reported that the bacterium generates toxins capable of directly affecting PMN and macrophages and, as a consequence, their function is altered. Additionally, the recruitment of M1 macrophages to the site of infection was affected by the lack of IFNγ. On the other hand, the bacterium generates damage at the level of the microvasculature and, as a consequence, affects the migration of the phagocytes into the damaged muscle. The alteration of the influx of the different inflammatory cells leads to a deficient cell debris removal of The above observations, coupled with the release of factors that stimulate the overproduction of collagen such as TGFβ1, result in poor muscle regeneration, characterized by fibrosis and small regenerative fibers.

Although muscle regeneration is highly efficient in many clinical and experimental models, provided basic requirements are fulfilled, muscle regeneration after myonecrosis induced by a sublethal inoculum of *C. perfringens* is deficient probably due to altered events in the initial phases following injury, which are critical and influence the overall outcome of the regeneration process. The inflammatory response induced by *C. perfringens* is characterized by alterations in the early influx of inflammatory cells, mainly PMN, macrophages M1 and M2 to the site of infection. These cells remain in the muscle for prolonged periods of time and are likely to be functionally impaired. In the case of M1 macrophages, their limited recruitment is possibly due to the low levels of INFγ produced. Our findings highlight several aspects of the regenerative muscle response which are affected in the experimental model of gas gangrene used. Understanding the mechanisms of the inflammatory and regenerative response in the muscle after infection by *C. perfringens* could be crucial for understanding the bases behind such poor regenerative response and for devising innovative therapeutic strategies for this drastic muscular pathology.

**Author contributions** Conceptualized, supervised and obtained funds for the project: MF-D; Conceived and designed experiments: MF-D, AA-G, AMZ-P, CS; Conducted experiments: AMZ-P, MF-D, CS; Analyzed data: MF-D, AA-G, AMZ-P, CS, JMG. Wrote the paper: MF-D, AA-G, AMZ-P; Edited the paper: MF-D, AA-G, AMZ-P, CS, JMG.

## Notes

**Funding:** This study was funded by Vicerrectoria de Investigation, Universidad de Costa Rica (Project No 741-B8-135 to MF-D). The funder had no role in study design, data collection, and analysis, decision to publish, or preparation of the manuscipt.

